# RhoGEF Solo regulates cell density-dependent keratin remodeling and desmosome maturation

**DOI:** 10.64898/2026.11.04.764450

**Authors:** Ukyo Kawasaki, Riku Maruta, Kensaku Mizuno, Shuhei Chiba, Kazumasa Ohashi

## Abstract

Desmosomes (DSMs) are specialized cell-cell adhesion structures that anchor keratin intermediate filaments and play a crucial role in maintaining the mechanical integrity of epithelial tissues. Keratin filaments associate with DSMs in two characteristic patterns: radial spokes extending from DSMs into the cytoplasmic keratin network and circumferential rims connecting neighboring DSMs. Solo is a Rho-guanine nucleotide exchange factor (RhoGEF) involved in cellular responses to mechanical stress and keratin network reorganization. Here, we investigated the role of Solo in regulating keratin-DSM interactions in response to cell density. The number of radial keratin spokes at cell-cell adhesion sites was markedly reduced at high cell density, concomitant with increased Solo localization at these sites. Conversely, *Solo* knockout (KO) increased the number of radial keratin spokes. Expression of wild-type Solo, but not a GEF-inactive Solo mutant, rescued the increase in radial keratin spokes in confluent *Solo*-KO cells, indicating that Solo RhoGEF activity is required for this regulation. Inhibition of actin polymerization or ROCK activity also suppressed the reduction in radial spoke density. Furthermore, expansion microscopy and electron microscopy revealed that DSMs in *Solo*-KO cells remained small and immature even under confluent conditions. Solo was also partially required for the localization of plakoglobin (PG), a core DSM protein, whereas *PG*-KO markedly increased the number of radial keratin spokes. These findings suggest that increased cell density promotes DSM maturation accompanied by a reduction in radial keratin spokes, and Solo contributes to both processes, potentially through facilitating PG localization at DSMs and regulating Rho-ROCK-dependent actin cytoskeletal remodeling.

## Introduction

Epithelial cells form organized layers that constitute higher-order tissues and organs. Within these multicellular structures, individual cells sense the state of neighboring cells through cell-cell adhesions and respond appropriately by regulating tissue homeostasis, cell morphology, differentiation, proliferation, and cell death (Garcia *et al*., 2018; Macara *et al*., 2014). These processes require coordinated cell movements and the maintenance of the mechanical integrity of epithelial sheets, both of which depend on dynamic cytoskeletal reorganization (Angulo-Urarte *et al*., 2020; Pandya *et al*., 2017). In particular, intercellular adhesion complexes and their associated cytoskeletal structures play critical roles in regulating collective cell motility, cell polarity, and the mechanical strength of epithelial sheets (Baum and Georgiou, 2011; Huber *et al*., 2015).

The actin cytoskeleton is a fundamental component of the cellular cytoskeletal network. Through interactions with myosin II, actin filaments generate contractile forces that regulate cell motility and morphological changes (Blanchoin *et al*., 2014). In epithelial cells, actin filaments also provide structural support for cell-cell junctions, including adherens junctions (AJs) and tight junctions (TJs), and contribute to the sensing and generation of intercellular mechanical forces (Lecuit *et al*., 2011). In vertebrates, cytoplasmic intermediate filaments (IFs) provide additional mechanical strength and structural integrity to cells (Broussard *et al*., 2020; Coulombe and Omary, 2002). In epithelial tissues, keratin IFs form extensive networks that are anchored to desmosomes (DSMs) at cell-cell adhesion sites and to hemidesmosomes at cell-substrate adhesion sites (Broussard et al., 2020). These keratin networks are connected to the cell membrane through plakin family proteins and to the nucleus through the linker of nucleoskeleton and cytoskeleton (LINC) complex. By spanning the cytoplasm, keratin IF networks contribute to cellular mechanical integrity (Garcia et al., 2018; Green *et al*., 2019; Zimmer and Kowalczyk, 2024).

Although keratin IFs form complex intracellular networks, their organization at cell-cell adhesion sites is particularly distinctive. Keratin IFs associated with DSMs form two characteristic structures: radial, spoke-like structures (hereafter referred to as “spokes”) and circumferential, rim-like structures (hereafter referred to as “rims”) (Quinlan *et al*., 2017). Keratin IFs associated with DSMs contribute to DSM maturation, the barrier function of cell-cell adhesions, and the mechanical integrity of epithelial tissues. Furthermore, actin cytoskeletal remodeling at cell-cell adhesion sites has been implicated in the regulation of keratin IF organization and DSM maturation (Janssen and Huveneers, 2024). Keratin IFs and the actin cytoskeleton are thought to regulate each other’s organization, thereby contributing to the maintenance of epithelial sheet integrity. However, the physiological significance of distinct keratin IF architectures at cell-cell adhesion sites and the molecular mechanisms governing their assembly and remodeling remain largely unknown.

We previously identified Solo as a Rho-guanine nucleotide exchange factor (RhoGEF) required for the alignment of vascular endothelial cells in response to cyclic stretching (Abiko *et al*., 2015). We subsequently identified keratin 8 (KRT8)/keratin 18 (KRT18) IFs as Solo-binding proteins and demonstrated that Solo contributes to the proper organization of KRT8/18 networks in Madin-Darby canine kidney (MDCK) cells (Fujiwara *et al*., 2016). Furthermore, we showed that Solo is involved in cellular responses to tensile forces applied to focal adhesions and that this response depends on the interaction between Solo and keratin IFs (Fujiwara *et al*., 2019). However, the specific keratin IF structures regulated by Solo remained unclear.

In this study, we established *Solo*-knockout (KO) MDCK cells and investigated the role of Solo in keratin IF organization at cell-cell adhesion sites. We found that Solo promotes the cell density-dependent remodeling of radial keratin IF spokes and contributes to DSM maturation. Our findings reveal a role for Solo in coordinating keratin IF organization and DSM maturation in response to changes in cell density.

## Materials & Methods

### Reagents and antibodies

Y-27632 (257-00511) was purchased from Wako Pure Chemical (Osaka, Japan). Latrunculin A (428021) and blebbistatin (203391) were purchased from Merck. Rabbit polyclonal antibodies against the following proteins were used: GFP (598, MBL), desmoglein 2 (DSG2) (21880-1-AP, Proteintech), wide-spectrum cytokeratin (ab9377, Abcam), and Solo, which was generated as described previously (Abiko et al., 2015). Mouse monoclonal antibodies against the following proteins were used: desmoplakin (DSP) (651109, Progen), b-catenin (610154, BD Biosciences), plakoglobin (610253, BD Biosciences), keratin 18 (KRT18) (61028, Progen), LARG (sc-25638, Santa Cruz), PDZ-RhoGEF (ab110059, Abcam) and a-tubulin (B-5-1-2, Abcam). A rat monoclonal antibody against keratin 8 (KRT8) (MABT329, Sigma-Aldrich) was used. Alexa Fluor 488-, Alexa Fluor 568-, and Alexa Fluor 633-conjugated goat anti-mouse IgG (A11029, A11031, and A21052, respectively), Alexa Fluor 488-conjugated goat anti-rabbit IgG (A11034), and Alexa Fluor 568-conjugated goat anti-rat IgG (A11077) were purchased from Thermo Fisher Scientific. Horseradish peroxidase (HRP)-conjugated sheep anti-mouse IgG (NA931) and donkey anti-rabbit IgG (NA934) were purchased from GE Healthcare.

### Cell culture

MDCK and GP2-293 cells were obtained from Riken BRD (Japan). Cells were cultured in Dulbecco’s modified Eagle’s medium (DMEM; 044-29765, Wako) supplemented with 10% fetal bovine serum (FBS) at 37°C in 5% CO_2_. MDCK cells stably expressing YFP-Solo were generated as described previously (Isozaki *et al*., 2020). To establish subconfluent and confluent conditions, 1.5 × 10⁵ and 7.5 × 10⁵ MDCK cells, respectively, were seeded per well of a 12-well plate and cultured for 2 days. Alternatively, the number of cells seeded and the culture period were adjusted to obtain mean cell areas of approximately 500 μm² and 150 μm² under subconfluent and confluent conditions, respectively.

### Plasmid construction

The expression plasmid encoding YFP-tagged Solo in the YFP-C1 vector was constructed as described previously (Abiko et al., 2015). The GEF-inactive Solo mutant, Solo (YALELE), was generated using a site-directed mutagenesis kit (Takara, Japan). Retroviral expression plasmids encoding EGFP-Solo and EGFP-Solo (YALELE) were constructed by inserting PCR-amplified cDNAs encoding EGFP-Solo or EGFP-Solo (YALELE) into the pRetroQ vector (Takara).

### Establishment of KO cell lines

KO cell lines were established using a homology-independent DNA repair method, as described previously (Katoh *et al*., 2017). Single-guide RNA (sgRNA) sequences targeting the canine *ARHGEF40* (Solo), *ARHGEF11* (PDZ-RhoGEF, PRG), *ARHGEF12* (leukemia-associated RhoGEF, LARG), or JUP (plakoglobin, PG) gene were designed using CRISPOR (https://crispor.gi.ucsc.edu). The sgRNA sequences are listed in Table 1. Double-stranded oligonucleotides corresponding to the target sequences were inserted into the all-in-one sgRNA expression plasmid pHiFiCAS9 (Addgene plasmid 162277; Fujisawa *et al*., 2021) and the donor plasmid pDonor-tBFP-NLS-Neo (Universal, Addgene plasmid #80767). MDCK cells cultured in 12-well plates were cotransfected with the sgRNA expression plasmid and the donor plasmid using Lipofectamine 2000 transfection reagent. Transfected cells were cultured and selected using G418 (750 µg/mL), and single colonies expressing tBFP were isolated. Genomic DNA was extracted from the isolated clones and subjected to PCR using KOD Fx Neo DNA polymerase (Toyobo, Japan) and three sets of primers designed to distinguish among three possible states of integration of the donor knock-in vector: forward integration, reverse integration, and no integration accompanied by a small insertion or deletion (indel). Disruption of both alleles of *Solo*, *LARG*, *PRG*, or *PG* genes was confirmed by direct sequencing of the PCR products.

**Table 1.**
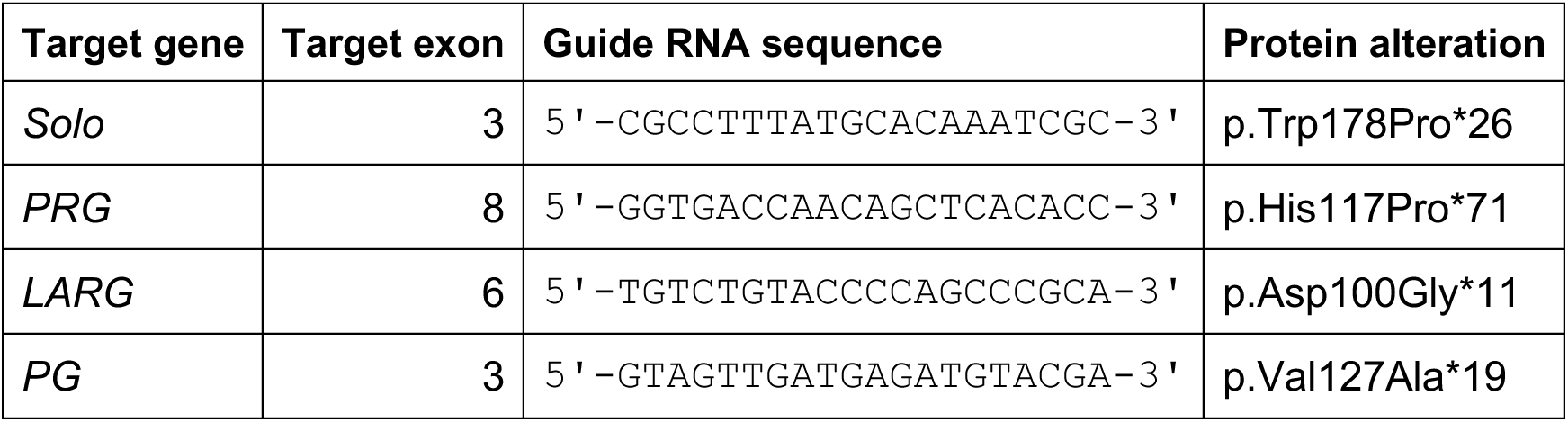
Target genes, guide RNA sequences, and predicted protein alterations.

### Retrovirus production

Retroviruses for expressing EGFP, EGFP-Solo, or the GEF-inactive Solo mutant [Solo (YALELE)] were produced by cotransfecting GP2-293 cells with the corresponding pRetroQ expression plasmids and the pMD2.G packaging plasmid using Lipofectamine 3000 transfection reagent (Thermo Fisher Scientific). The culture medium was replaced 4 h after transfection, and the cells were cultured for an additional 2 days. The virus-containing culture medium was collected at 24 h and 48 h after transfection and passed through a 0.45-µm filter.

### Establishment of stable cell lines

MDCK cells stably expressing YFP-Solo were established previously (Isozaki *et al*., 2020). *Solo*-KO cells stably expressing EGFP-Solo or EGFP-Solo (YALELE) mutant were generated by retroviral infection. *Solo*-KO cells were infected with the respective retrovirus in growth medium containing polybrene (8 µg/mL) for 24 h, followed by selection with puromycin (2 µg/mL). Several individual clones were isolated from the selected cells, and expression of the EGFP-tagged proteins was confirmed by fluorescence microscopy and western blotting using an anti-GFP antibody.

### Fractionation of KRT8/18

Total cell lysates were prepared by solubilizing MDCK cells cultured under subconfluent or confluent conditions in lysis buffer (62.5 mM Tris-HCl, pH 6.8, 2% SDS, 5% β-mercaptoethanol, and 5% sucrose). Keratin-soluble and keratin-insoluble fractions were prepared according to a previously described procedure (Busch *et al*., 2012). In brief, MDCK cells were scraped into PBS containing 1% Nonidet P-40, 10 mM EDTA, and 1× protease inhibitor cocktail (Nacalai Tesque, Japan) and incubated for 15 min at 4°C. The samples were then centrifuged at 16,000 × g for 15 min at 4°C. The supernatant and precipitate were separately denatured and solubilized in Laemmli sample buffer and designated the soluble and insoluble fractions, respectively. These fractions were separated by SDS-PAGE and analyzed by western blotting.

### Ca^2+^-switch assay

Control MDCK and *Solo*-KO MDCK cells were cultured on 15-mm-diameter coverslips (C015001 Matsunami, Japan) in 12-well plates under subconfluent or confluent conditions. To deplete Ca^2+^ from the culture medium, the cells were washed with phosphate-buffered saline (PBS) and cultured for 12 h in Ca^2+^- and L-glutamine-free DMEM (21068028, Gibco), supplemented with 10% Ca^2+^-depleted FBS and L-glutamine. The cells were then returned to normal culture medium, and this time point was designated time 0. Ca^2+^ depleted FBS was prepared by incubating FBS with Chelex 100 resin (142-2832, Bio-Rad) for 12 h, followed by filtration to remove the Ca^2+^-adsorbed resin.

### Immunofluorescence staining and image analysis

MDCK cells cultured on 15-mm-diameter coverslips were fixed with 4% paraformaldehyde (PFA) in PBS for 15 min at 37°C or cold methanol for 10 min at −20°C. The cells were then permeabilized with 0.1% Triton X-100 in PBS for 5 min at room temperature. After washing with PBS, the cells were blocked with 2% FBS in PBS and then incubated overnight at 4°C with primary antibodies diluted in 2% FBS in PBS. The cells were washed with PBS and incubated with secondary antibodies diluted in 2% FBS in PBS for 60 min, followed by washing with PBS. Coverslips were mounted on slide glasses using Mowiol (Calbiochem). Fluorescence images were obtained using a THUNDER Ready fluorescence microscope (Leica Microsystems) equipped with PL Apo 100× and 63× oil objective lenses (1.4 and 1.3 NA, respectively), and an ORCA-Flash4.0 digital CMOS camera (Hamamatsu Photonics, Japan). Images were analyzed using Fiji software (Fiji, an open-source platform for biological-image analysis). Fluorescence intensities of YFP-Solo, PG, and keratin at cell-cell adhesion sites were measured using Z-stack images acquired at 0.21 μm intervals. The Z-stack images were processed using maximum-intensity projection. Cell-cell adhesion sites were identified based on β-catenin or DSP signals, and the corresponding junctional lines were traced. Fluorescence intensities of YFP-Solo, keratin and PG at the cell-cell adhesion sites were measured within a 10-pixel-wide region centered on each traced line. The fluorescence intensity at cell-cell adhesion sites was normalized to that in the cytoplasm. To quantify keratin spoke density at cell-cell adhesion sites, maximum-intensity projection images were generated from seven Z-stack images acquired at 0.21 μm intervals, comprising the focal plane containing DSP signals and three optical sections above and below that plane. The number of keratin filaments extending from DSP-positive sites toward the cytoplasm (keratin spokes) at cell-cell adhesion sites was manually counted in the processed images. Spoke density was expressed as the number of spokes per 1 μm of cell-cell adhesion. Desmosome length was measured as previously described (Beggs *et al*., 2021; Beggs *et al*., 2022). Desmosomes were manually selected from a single z-plane of the ExM images or from TEM images. For each desmosome, a line was drawn along the DSP signal parallel to the plasma membrane, and the length of this line was defined as the desmosome length (Figure 5D).

### Western blotting

Total cellular proteins were separated by SDS-PAGE and transferred to a polyvinylidene difluoride (PVDF) membrane (Immobilon, MILLIPORE). The membranes were blocked with 5% nonfat dry milk in PBS containing 0.05% Tween20 (PBS-T) for 1 h at room temperature. After washing with PBS-T, the membranes were incubated with primary antibodies in 1% nonfat dry milk in PBS-T, and then with horseradish peroxidase (HRP)-conjugated secondary antibodies. Chemiluminescence signals were generated using Immobilon Western Chemiluminescent HRP Substrate or Immobilon ECL Ultra Western HRP Substrate (Millipore) and detected using a Chemidoc Touch imaging system (Bio-Rad).

### Uni-ExM

Uni-ExM samples were prepared with minor modifications to the original protocol previously described (Chen *et al*., 2015; Cui *et al*., 2023). Cells were fixed and stained with anti-DSG2 and anti-DSP antibodies, as described above. Before the crosslinking reaction, the stained cells were pre-incubated twice with 100 mM NaHCO_3_ (pH 8.5) for 15 min each. The cells were then incubated overnight at room temperature in 100 mM NaHCO_3_ (pH 8.5) containing 0.04% (w/v) glycidyl methacrylate. After the anchoring reaction, the cells were washed three times with PBS, and then the standard expansion procedures, including gelation, protein digestion, and expansion, were performed. For gelation, the cells were incubated with a gelation solution prepared by mixing monomer solution (MS; 8.6% (w/v) sodium acrylate, 2.5% (w/v) acrylamide, 0.15% N,N’-methylenebisacrylamide, and 2 M NaCl in PBS), 10% ammonium persulfate (APS), N,N,N’,N’-tetramethylethylenediamine (TEMED) (cat. 205-06313, Fujifilm, Japan), and water at a ratio of 470:10:1:19. A 45-µl drop of the gelation solution was placed on a Petri dish lined with Parafilm (Fisher Scientific) and cooled on ice. The coverslip (15 mm in diameter) bearing the cells was placed on the drop and incubated for 15 min on ice, followed by incubation for 60 min at 37°C. The coverslip with the attached gel was then transferred to a 12-well plate and immersed in digestion solution (50 mM Tris, 1 mM EDTA, 0.5% Triton X-100, 0.8 M guanidine HCl, and 6 units/mL proteinase K) for 45 min at 37°C. After digestion, the solution was removed, and the gel was washed twice with PBS for 30 min each. The gel was then immersed in water for 5–30 min. This step was repeated four times until the gel was fully expanded. DAPI-containing water was used for the second water exchange to stain the nuclei. After expansion, the gels were transferred to glass-bottom dishes (coverglass thickness, 0.16–0.19 mm; Matsunami, Japan), with the cell-containing surface facing the glass bottom and the basal surface facing the objective. The gels were immobilized with 2% agarose solution to prevent movement during imaging.

### TEM analysis

Cells were cultured in 3.5-cm dishes until confluence. The cells were fixed at 4°C for 30 min in 0.1 M phosphate buffer (pH 7.4) containing 2% paraformaldehyde and 2% glutaraldehyde, followed by postfixation in 0.1 M phosphate buffer (pH 7.4) containing 2% glutaraldehyde. The samples were submitted to Tokai Electron Microscopy for TEM imaging.

### Dispase assay

Cells were cultured to confluence in 24-well culture plates. After washing with PBS containing 1 mM CaCl_2_, the cells were incubated with dispase II (1000 PU/mL; Fujifilm) for 1 h. The released cell monolayers were subjected to mechanical stress by vortexing for 30 s, and the resulting fragments were counted manually.

### Statistical analyses

Data are presented as the mean ± SD of at least three independent experiments. Statistical analyses were performed using Prism 6 (GraphPad Software, La Jolla, CA, USA). *P* values were calculated using Student’s *t*-test or one-way ANOVA, followed by Dunnett’s or Sidak’s multiple-comparisons test.

## Results

### 1. Increased cell density facilitates Solo localization at cell-cell adhesion sites

We previously showed that Solo localizes in a punctate pattern at cell-cell adhesion sites in MDCK cells (Isozaki *et al*., 2020). To investigate whether cell density affects Solo localization, we established MDCK cells constitutively expressing YFP-Solo and examined its localization under subconfluent (low-cell-density) and confluent (high-cell-density) conditions. YFP-Solo localization at cell-cell adhesion sites was significantly increased under confluent conditions, compared with subconfluent conditions (Figure 1A, B). These results indicate that increased cell density promotes Solo localization at cell-cell adhesion sites, suggesting that Solo may contribute to cell density-dependent remodeling of keratin structures at these sites.

**Figure 1.**
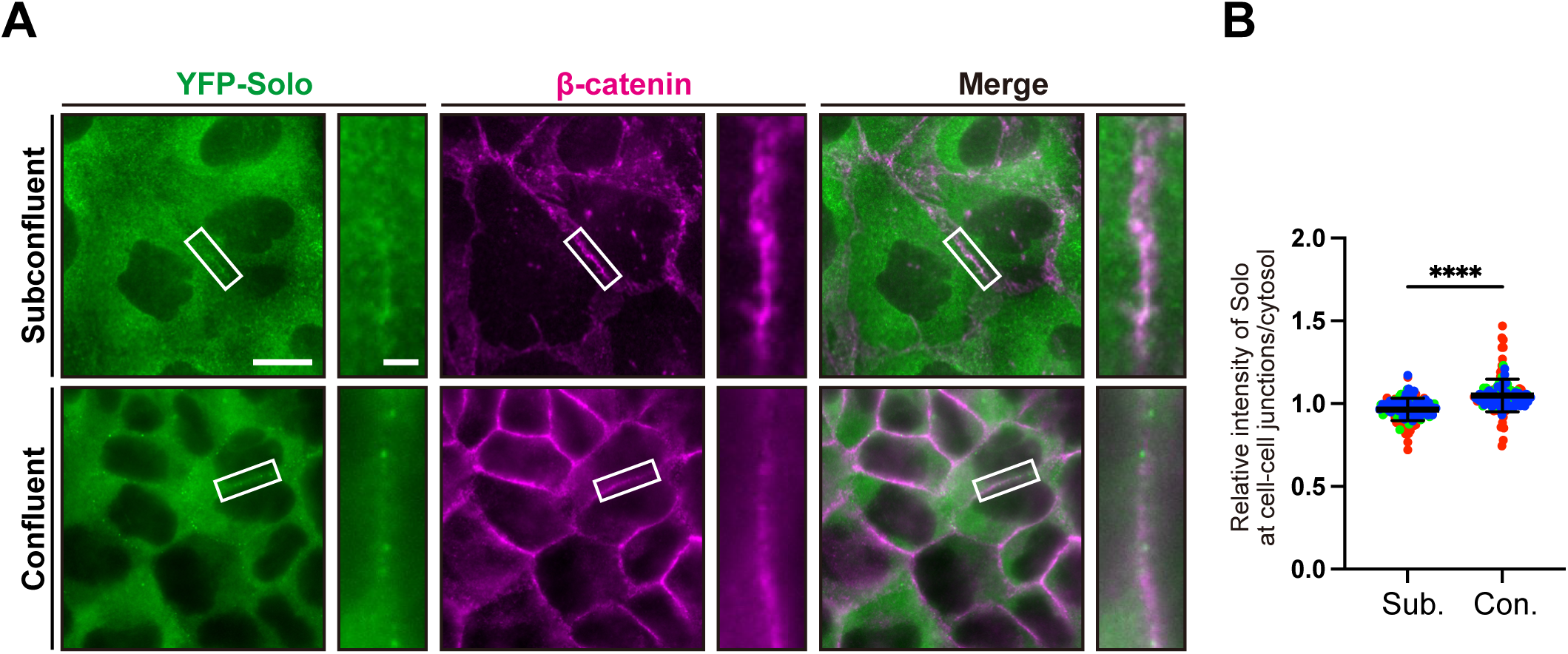
Solo accumulates at cell-cell adhesion sites under high-density confluent culture conditions. (A) Representative images of YFP-Solo (green) in MDCK cells stably expressing YFP-Solo cultured under subconfluent (upper panels) or confluent (lower panels) conditions. Cells were fixed and stained with an anti-β-catenin antibody to visualize cell-cell adhesion sites (magenta). The right panels show magnified views of the boxed regions in the corresponding left panels. Scale bars, 10 µm (magnified images, 2 µm). (B) Quantification of the relative intensity of YFP-Solo at cell-cell adhesion sites under subconfluent (Sub.) and confluent (Con.) conditions. Data are presented as the mean ± SD from three independent experiments, with 42-56 cell-cell junctions analyzed per experiment. ****, *p* < 0.0001 (Student’s t-test).

### 2. Radial keratin spokes are reduced under confluent conditions

Because Solo localization at cell-cell adhesion sites varied with cell density, we next investigated whether keratin IF organization at these sites also changes in a cell density-dependent manner. Keratin IFs adopt distinct structural configurations at cell-cell adhesion sites, including radial spoke and circumferential rim structures (Figure 2A; Quinlan et al., 2017). We first confirmed differences in cell density between subconfluent and confluent conditions by measuring the mean cell area (Figure 2B). Numerous radial keratin spokes were observed under subconfluent conditions, whereas their number was markedly reduced under confluent conditions (Figure 2A, C). In contrast, the relative fluorescence intensity of keratin IFs at cell-cell adhesion sites, normalized to that in the cytoplasmic region, increased under confluent conditions (Figure 2D). These findings indicate that increased cell density is associated with a reduction in radial keratin spokes despite an increase in the relative accumulation of keratin IFs at cell-cell adhesion sites. In subsequent experiments, the cell densities were adjusted to achieve the mean cell areas shown in Figure 2B.

**Figure 2.**
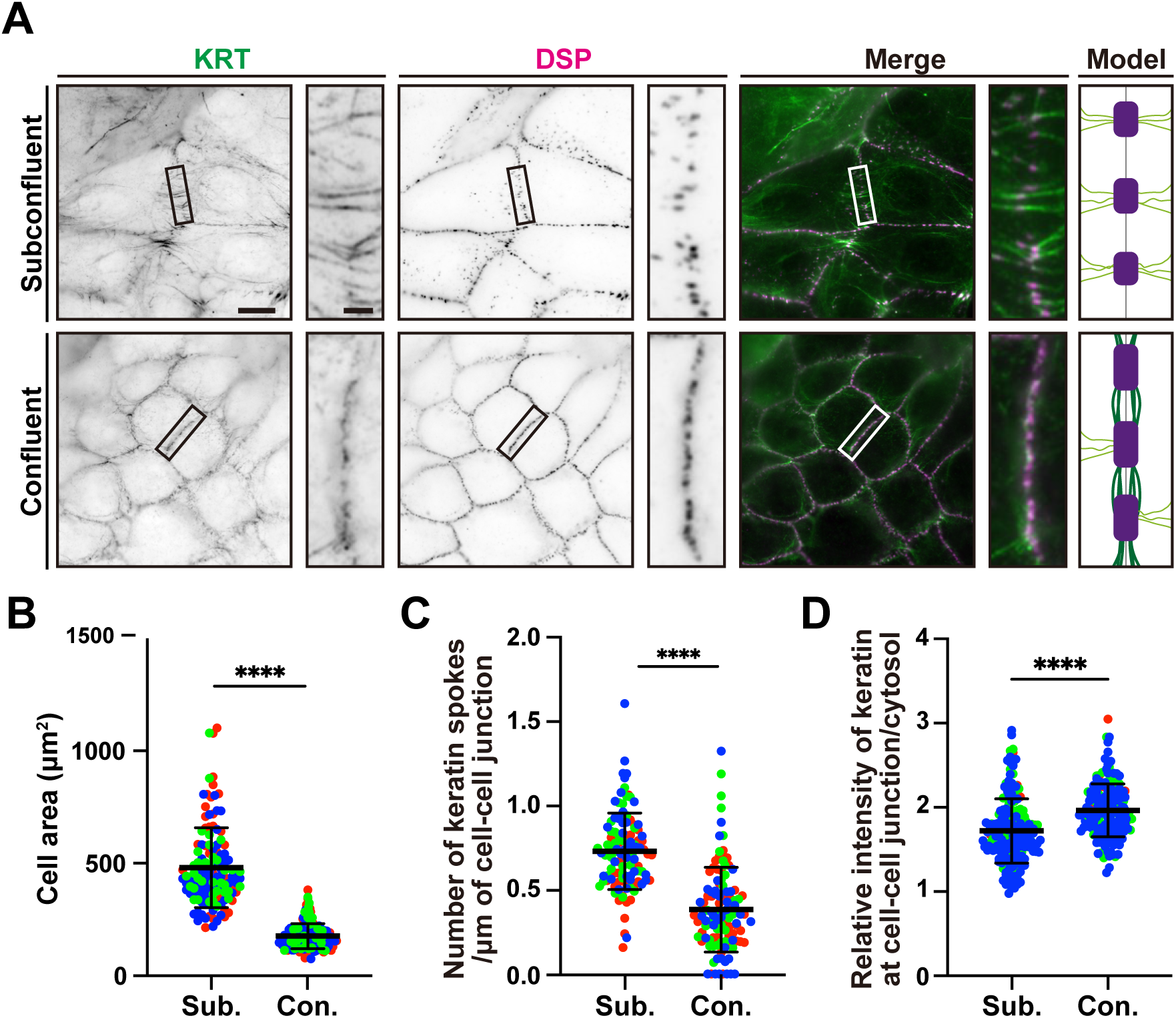
Radial keratin spokes and circumferential keratin rims are regulated in a cell density-dependent manner. (A) Representative images of keratin filaments (KRT; green) and desmoplakin (DSP; magenta), accompanied by schematics illustrating keratin filament organization in MDCK cells cultured under subconfluent (upper) and confluent (lower) conditions. Cells were fixed and stained with anti-desmoplakin (DSP) and anti-pan-keratin (KRT) antibodies. The right panels show magnified views of the boxed regions in the corresponding left panels. Scale bars, 10 µm (magnified images, 2 µm). In the schematics, desmosomes are depicted in purple, radial keratin spokes in light green, and circumferential rim-like keratin filaments in dark green. (B) Quantification of cell area under subconfluent (Sub.) and confluent (Con.) conditions. Data are presented as mean ± SD from three independent experiments (39-51 cells per experiment). ****, *p* < 0.0001 (Student’s t-test). (C) Quantitative analysis of spoke density at cell-cell adhesion sites under subconfluent and confluent conditions. Data are presented as the mean ± SD from three independent experiments, with 30-40 cell-cell junctions analyzed per experiment. ****, *p* < 0.0001 (Student’s t-test). (D) Relative fluorescence intensity of keratin filaments at cell-cell adhesion sites compared with that in the cytoplasm under subconfluent and confluent conditions. Data are presented as the mean ± SD from three independent experiments, with 52-67 cell-cell junctions analyzed per experiment. ****, *p* < 0.0001 (Student’s t-test).

### 3. Solo is required for the reduction of radial keratin spokes at cell-cell adhesion sites under confluent conditions

To elucidate the role of Solo in regulating keratin IF organization at cell-cell adhesion sites, we established *Solo*-KO MDCK (*Solo*-KO) cells (Supplemental Figure S1A, B). The absence of Solo protein in *Solo*-KO cells was confirmed by immunoblotting of cell lysates using an anti-Solo antibody (Supplemental Figure S1B). Because KRT8 and KRT18 are the major cytoplasmic keratins in MDCK cells, we examined whether Solo depletion affects KRT8 and KRT18 expression levels or the KRT8/18 soluble-to-insoluble ratio. Neither parameter differed between *Solo*-KO and parental MDCK (control) cells (Supplemental Figure S1C).

We next examined the effects of Solo depletion on radial keratin spoke density at cell-cell adhesion sites. Spoke density was significantly higher in *Solo*-KO cells than in control cells under both subconfluent (Figure 3A, B) and confluent conditions (Figure 3C, D). Notably, the difference in spoke density between control and *Solo*-KO cells was particularly pronounced under confluent conditions (Figure 3C, D). To determine whether the GEF activity of Solo is required for the reduction in spoke density under confluent conditions, we performed rescue experiments using *Solo*-KO cells expressing EGFP alone, EGFP-tagged wild-type Solo (WT), or the GEF-inactive mutant Solo (YALELE). Under confluent conditions, expression of EGFP-Solo (WT), but not EGFP alone, reduced spoke density in *Solo*-KO cells to levels comparable to those observed in control cells (Figure 3C, D). Meanwhile, expression of the Solo (YALELE) mutant further increased spoke density in Solo-KO cells. Immunoblotting confirmed the expression of EGFP alone and EGFP-Solo (WT) (Supplemental Figure S1D). These results indicate that Solo is required for the reduction of radial keratin spoke density under confluent conditions and that its GEF activity is necessary for this regulation.

**Figure 3.**
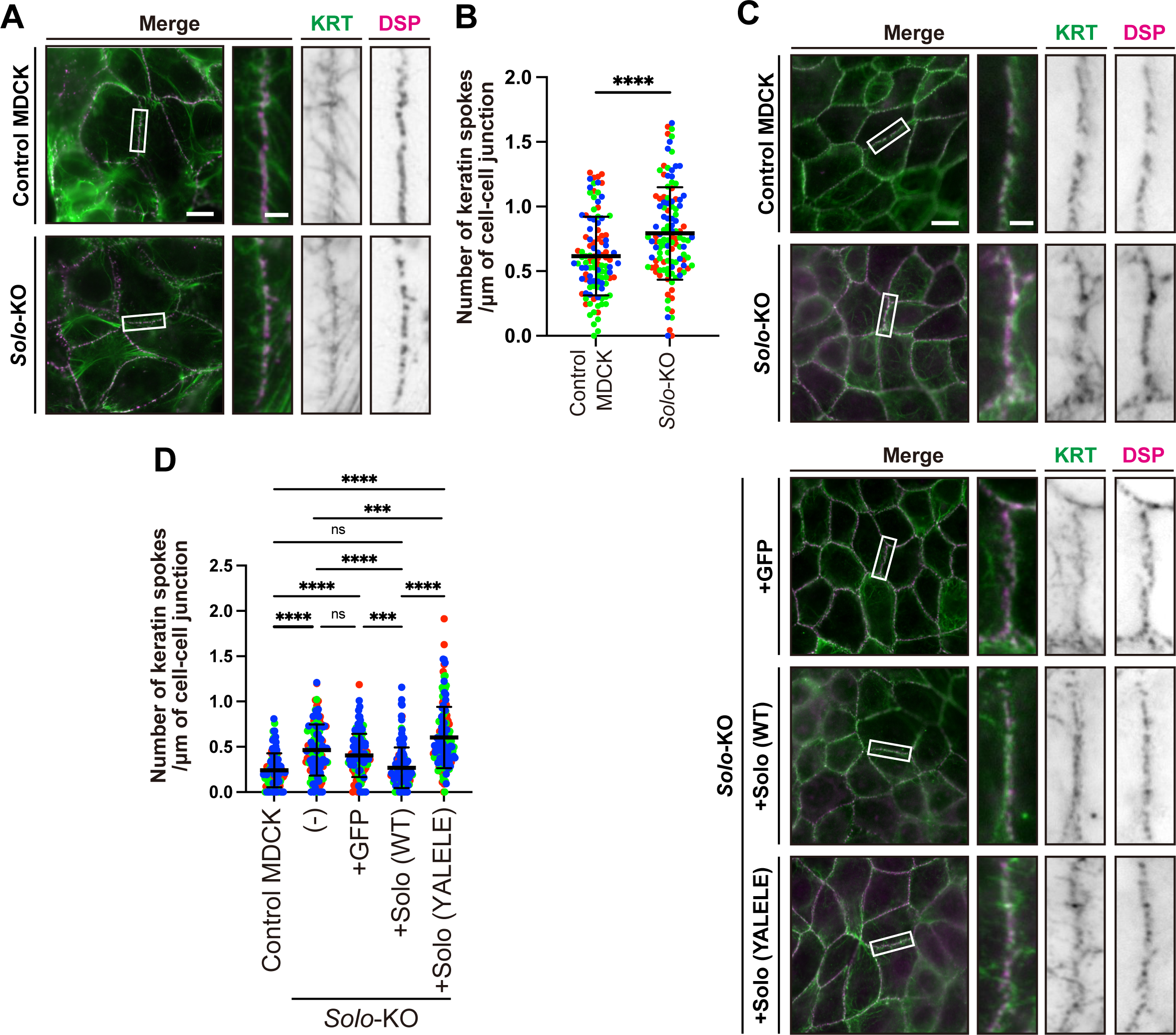
Solo regulates the density-dependent reduction of keratin spokes. (A) Representative images of keratin networks of parental (Control) MDCK and *Solo*-KO cells under subconfluent conditions. Cells were fixed and stained as described in Fig. 2A (KRT; green, DSP; magenta). The right panels show magnified views of the boxed regions at cell-cell adhesion sites in the corresponding left panels. Scale bars, 10 µm (magnified images, 2 µm). (B) Quantitative analysis of the density of spokes at cell-cell adhesion sites shown in (A). Data are presented as the mean ± SD from three independent experiments, with 33-50 cell-cell junctions analyzed per experiment. ****, *p* < 0.0001 (Student’s t-test). (C) Representative images of keratin networks in control MDCK cells and *Solo*-KO cells expressing EGFP, EGFP-Solo (WT), or EGFP-Solo (YALELE) mutant under confluent conditions. Cells were fixed and stained as described in (A). (D) Quantification of the density of keratin spokes at cell-cell adhesion sites, as shown in (C). Data are presented as the mean ± SD from three independent experiments, with 31-59 cell-cell junctions analyzed per experiment. ***, *p* < 0.001; ****, *p* < 0.0001; ns, not significant (one-way ANOVA with Sidak’s multiple comparisons test).

### 4. Reduction in radial keratin spoke density depends on cell density rather than the duration of cell-cell contact

We hypothesized that radial keratin spoke density changes as cell density increases. To determine whether the reduction in spoke density depends on the duration of cell-cell contact or on cell density, we performed a Ca²⁺-switch assay (Figure 4). MDCK cells were seeded under subconfluent or confluent conditions and allowed to form cell-cell adhesions. Cell-cell adhesions were then disrupted by culturing the cells in low-Ca²⁺ medium for 12 h. Following Ca²⁺ restoration, cells were fixed at various time points over the subsequent 24 h, and keratin IFs were visualized to quantify spoke density at cell-cell adhesion sites (Figure 4A, B). Spoke density increased shortly after cell-cell contacts were re-established under both subconfluent and confluent conditions. Under subconfluent conditions, spoke density continued to increase throughout the 24-h period (Figure 4B). In contrast, under confluent conditions, spoke density initially increased, peaked approximately 6 h after Ca^2+^ restoration, and then gradually decreased over the remainder of the observation period (Figure 4B). These results indicate that the reduction in radial keratin spoke density is associated with cell density rather than simply with the duration of cell-cell contact. Because *Solo*-KO cells exhibited increased spoke density under confluent conditions, we next performed a Ca^2+^-switch assay using confluent *Solo*-KO cells. Spoke density in *Solo*-KO cells was comparable to that in control cells for up to 6 h after Ca^2+^ restoration. However, the subsequent reduction in spoke density was impaired in *Solo*-KO cells (Figure 4A, B). These findings indicate that Solo is required for the cell density-dependent reduction of radial keratin spokes at cell-cell adhesion sites.

**Figure 4.**
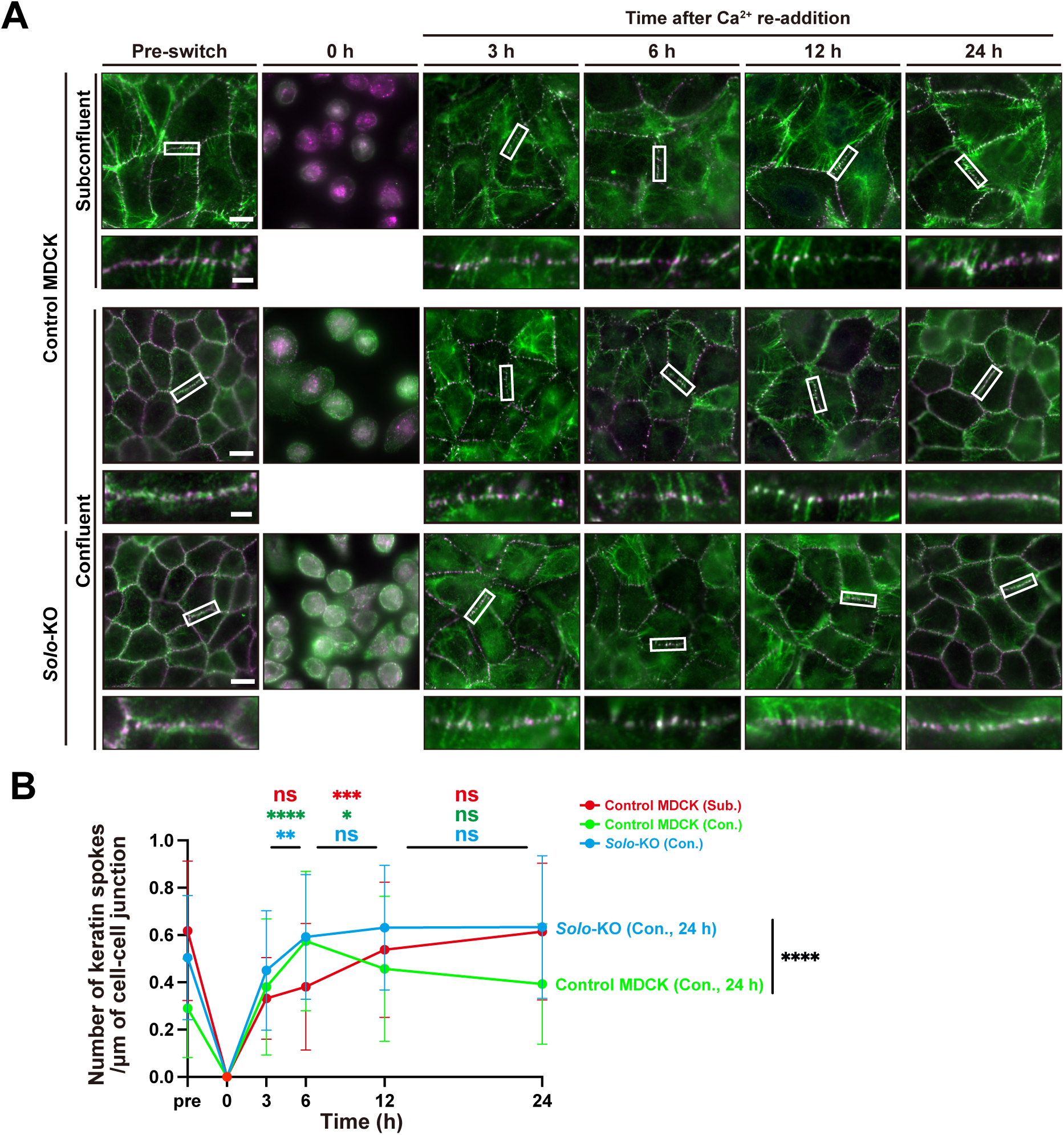
Radial keratin spokes are assembled during the early stages of cell-cell junction formation and gradually decrease under confluent conditions. (A) Representative images of the keratin organization in parental (Control) MDCK and *Solo*-KO cells during the Ca^2+^-switch assay under subconfluent (Control) and confluent (Control and *Solo*-KO cells) conditions. Cells were cultured under confluent or subconfluent conditions, fixed, and stained as described in Fig. 2A before and at the indicated times following the Ca^2+^ restoration. The lower panels show magnified views of the boxed regions in the corresponding upper panels. Scale bars, 10 µm (magnified images, 2 µm). (B) Quantitative analysis of the time-dependent changes in radial spoke density following the formation of cell-cell adhesions shown in (A). Data are presented as mean ± SD from three independent experiments, with 30 cell-cell junctions analyzed per experiment. Statistical comparisons were performed between 3 and 6 h, 6 and 12 h, and 12 and 24 h, using one-way ANOVA with Sidak’s multiple comparisons test. *, *p* < 0.05; **, *p* < 0.01; ***, *p* < 0.001; ****, *p* < 0.0001; ns, not significant. Spoke density was also compared between control and *Solo*-KO cells after 24 h under confluent conditions, using Student’s t-test. ****, *p* < 0.0001.

### 5. Solo is involved in desmosome maturation

DSMs are known to increase in size through fusion with one another during maturation (Beggs et al., 2022; Gloushankova *et al*., 2003). We hypothesized that *Solo*-KO cells would fail to increase DSM size under confluent conditions and consequently retain a higher density of radial keratin spokes. To test this hypothesis, we analyzed DSM morphology in control and *Solo*-KO cells using expansion microscopy (ExM), a super-resolution imaging technique (Chen et al., 2015; Cui et al., 2023). ExM achieves enhanced spatial resolution by embedding biological samples in a swellable hydrogel that physically expands the samples approximately 4–4.5-fold. In our experiments, the gels expanded by 4.14 ± 0.05-fold (Figure 5A, B). Assuming a conventional lateral optical resolution of 250-300 nm, this expansion corresponds to an effective lateral resolution of approximately 60-70 nm. ExM imaging following immunostaining for desmoplakin (DSP) and desmoglein 2 (DSG2) revealed patch-like structures characteristic of DSMs along both sides of the cell membranes at cell-cell junctions. DSM length was quantified based on DSP staining. In control cells, DSM length along the cell-cell junctions was greater under confluent conditions than under subconfluent conditions (Figure 5C-E). In contrast, DSMs were significantly shorter in *Solo*-KO cells than in control cells under confluent conditions, with lengths comparable to those observed in control cells under subconfluent conditions (Figure 5C, E). To further investigate these differences, we examined cross-sections of cell-cell adhesion sites by transmission electron microscopy (TEM). DSMs appeared as electron-dense structures at cell-cell adhesion sites. Consistent with the ExM results, DSM length was significantly shorter in *Solo*-KO cells than in control cells under confluent conditions (Figure 5F, G). In control cells, keratin filaments running parallel to the cell membrane accumulated at DSMs and formed rim-like structures connecting adjacent DSMs, whereas relatively few keratin filaments were observed anchoring perpendicular to the DSMs. In contrast, *Solo*-KO cells exhibited a lower density of keratin filaments within the rim-like structures, while perpendicular keratin filaments were more prominent (Figure 5F). Taken together, these results suggest that Solo contributes to DSM maturation under confluent conditions. The shorter DSMs and altered keratin filament organization observed in *Solo*-KO cells further suggest that DSM maturation is associated with a shift in the keratin filament organization from prominent perpendicular (radial) keratin filament anchoring toward the formation of rim-like structures connecting adjacent DSMs.

**Figure 5.**
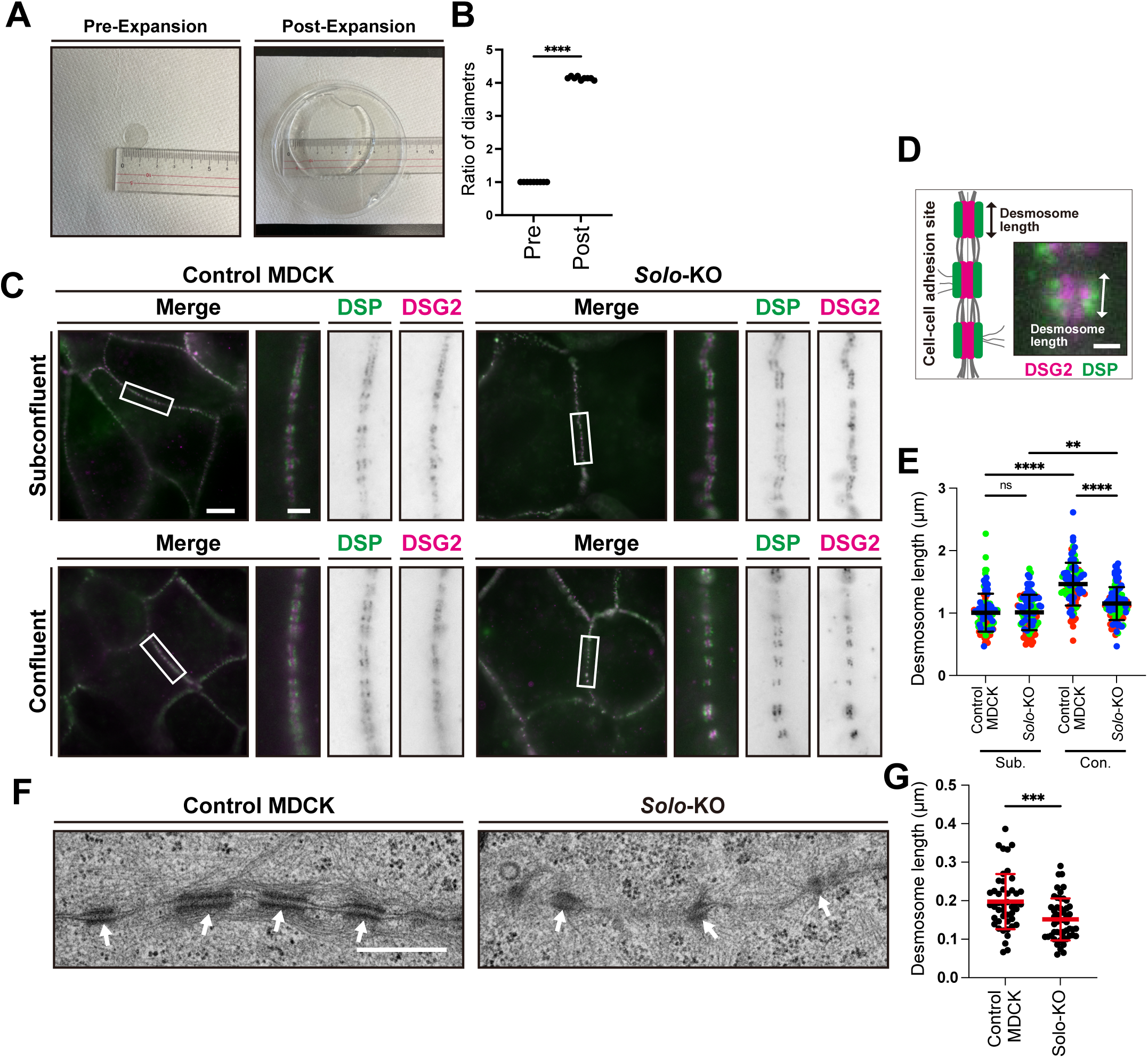
Solo is required for desmosome maturation. (A) Representative photographs of a hydrogel containing cross-linked and embedded MDCK cells, taken before and after expansion. (B) Quantification of the expansion ratio of the acrylamide gels, calculated from the gel diameters shown in (A). Data are presented as the mean ± SD from 9 independent experiments. ****, *p* < 0.0001 (Student’s t-test). (C) Representative ExM images of DSMs in control and *Solo*-KO cells cultured under subconfluent and confluent conditions. Cells were fixed and stained with anti-DSP (green) and anti-DSG2 (magenta) antibodies, followed by ExM treatment (see Methods). The right panels show magnified views of the boxed region in the corresponding left panels. Scale bars, 40 µm (magnified images, 8 µm), corresponding to an estimated actual size of 10 µm (magnified images, 2 µm). (D) Definition of DSM length. DSM length was defined as the length of the DSP signal parallel to the cell membrane. The schematic illustrates the cell–cell adhesion site with DSMs, and the right image shows a DSM visualized by immunofluorescence staining for DSG2 (magenta) and DSP (green) in an ExM image. Scale bars, 1 µm. (E) Quantification of DSM length shown in (C). Data are presented as the mean ± SD from three independent experiments, with 27-43 DSMs analyzed per experiment. **, *p* < 0.01; ****, *p* < 0.0001; ns, not significant (one-way ANOVA with Sidak’s multiple comparisons test). (F) Representative TEM micrographs of horizontal cross-sections of cell-cell adhesion sites in control and *Solo*-KO cells. Arrows indicate DSMs. Scale bars, 200 nm. (G) Quantitative analysis of DSM length shown in (F). Data are presented as the mean of 47 DSMs from control cells and 50 DSMs from *Solo*-KO cells. ***, *p* < 0.001 (Student’s t-test).

### 6. Reduction of radial keratin spokes under confluent conditions requires actin cytoskeletal remodeling through the Rho-ROCK pathway

We next investigated whether actin cytoskeletal remodeling contributes to the cell density-dependent reduction in radial keratin spokes. Previous studies have shown that pharmacological perturbation of the actin cytoskeleton using latrunculin A, an inhibitor of actin polymerization; Y-27632, an inhibitor of Rho-associated coiled-coil-containing protein kinase (ROCK); and blebbistatin, an inhibitor of actomyosin contractility, affects keratin spoke formation (Prechova *et al*., 2022). We therefore examined whether actin polymerization and/or actomyosin contractility is involved in the reduction of radial keratin spokes under confluent conditions using these reagents. Latrunculin A and Y-27632 suppressed the reduction in radial keratin spoke density under confluent conditions, whereas blebbistatin had no detectable effect, consistent with previous findings (Figure 6A, B; Prechova *et al*., 2022). As shown in Figure 3, Solo depletion impaired the reduction in radial keratin spoke density under confluent conditions. Expression of Solo (WT) rescued this phenotype in *Solo*-KO cells, whereas expression of the GEF-inactive Solo (YALELE) mutant further increased spoke density (Figure 3A–D). Together, these findings suggest that Solo GEF activity and actin cytoskeletal remodeling are required for the reduction in radial keratin spoke density under confluent conditions, implicating Solo and the RhoA-ROCK pathway in this process.

**Figure 6.**
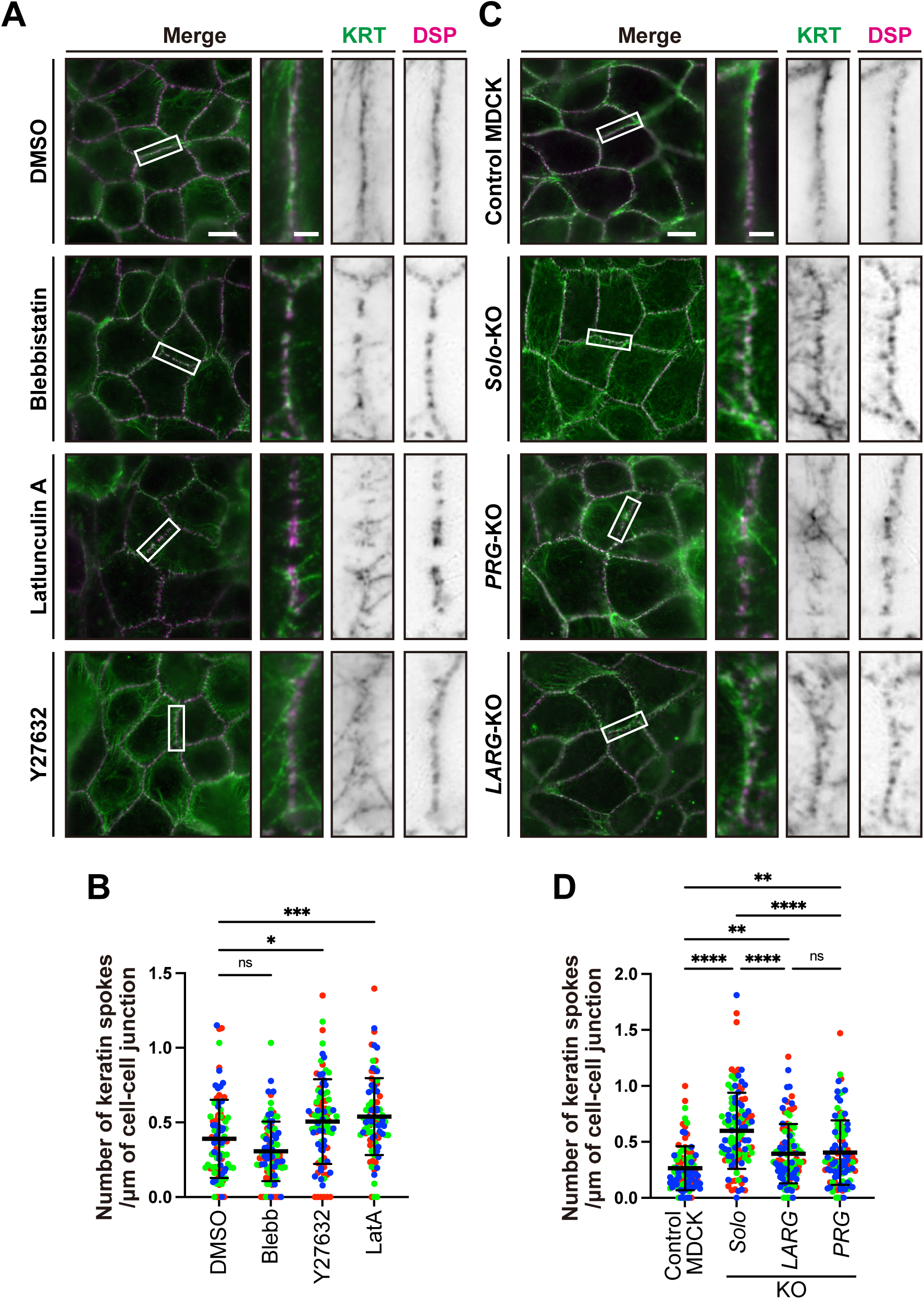
Density-dependent reduction of keratin spokes is regulated by actin cytoskeletal remodeling through the Rho/ROCK pathway. (A) Representative images of keratin networks of MDCK cells treated with blebbistatin (50 µM), latrunculin A (200 nM), or Y-27632 (10 µM) for 1 h under confluent conditions. Cells were fixed and stained with anti-pan-keratin (green) and anti-desmoplakin (DSP; magenta) antibodies. The right three panels show magnified views of the boxed regions in the corresponding left panels. Scale bars, 10 µm (right three panels, 2 µm). (B) Quantitative analysis of spoke density at cell-cell adhesion sites shown in (A). Data are presented as the mean ± SD from three independent experiments, with 30 cell-cell junctions analyzed per experiment. *, *p* < 0.05; ***, *p* < 0.001; ns, not significant (one-way ANOVA followed by Dunnett’s multiple comparisons test). (C) Representative images of keratin networks in control, *Solo*-KO, *PRG*-KO, and *LARG*-KO MDCK cells cultured under confluent conditions. Cells were fixed and stained with anti-DSP and anti-keratin antibodies. The right three panels show magnified views of the boxed regions in the corresponding left panels. Scale bars, 10 µm (right three panels, 2 µm). (D) Quantitative analysis of spoke density at cell-cell adhesion sites shown in (C). Data are presented as the mean ± SD from three independent experiments, with 30-42 cell-cell junctions analyzed per experiment. **, *p* < 0.01; ****, *p* < 0.0001; ns, not significant (one-way ANOVA with Sidak’s multiple comparisons test).

We previously reported that Solo promotes RhoA activation through the RhoA GEFs, PDZ-RhoGEF (PRG) and leukemia-associated Rho GEF (LARG) (Kunitomi *et al*., 2024; Kunitomi *et al*., 2026). We therefore investigated whether PRG and LARG are also involved in the reduction of radial keratin spoke density. We established *PRG*- and *LARG*-KO cells (Supplemental Figure S2A–D) and analyzed the effects of PRG or LARG depletion on spoke density at cell-cell adhesion sites. Both *PRG*-KO and *LARG*-KO cells exhibited significantly increased spoke density, although the extents of the increases were smaller than that observed in *Solo*-KO cells (Figure 6C, D). These findings suggest that PRG and LARG contribute to the regulation of radial keratin spoke density downstream of Solo and may have partially redundant roles in RhoA activation at cell-cell adhesion sites.

### 7. Solo regulates plakoglobin localization, and plakoglobin contributes to the reduction of radial keratin spoke density

We previously demonstrated that knockdown of Solo reduces the localization of plakoglobin (PG), a component of DSMs, at cell-cell adhesion sites (Isozaki et al., 2020). To further investigate the relationship between Solo and PG, we examined PG localization at cell-cell adhesion sites in control and *Solo*-KO cells. PG localization was partially but significantly reduced in *Solo*-KO cells compared with control cells (Figure 7A, B). We next investigated the role of PG in regulating radial keratin spoke formation by establishing *PG*-KO cells (Supplemental Figure S2E, F). Under confluent conditions, *PG*-KO cells exhibited a marked increase in radial keratin spoke density compared with control cells (Figure 7C, D). These findings suggest that PG contributes to the reduction of radial keratin spoke density at cell-cell adhesion sites under confluent conditions and that Solo may promote this process, at least in part, by facilitating PG localization at DSMs. PG has been reported to interact with desmoplakin at DSMs and to contribute to the formation and maintenance of DSM structures. Consistent with this role, loss of PG has been shown to substantially weaken epithelial cell-cell adhesion (Yin *et al*., 2005). To determine whether the increased radial keratin spoke density observed in *Solo*-KO cells was associated with altered mechanical integrity of the epithelial cell layer, we assessed the mechanical strength of cell layer using a dispase-based dissociation assay. *Solo*-KO cells did not exhibit a significant change in the number of cell layer fragments compared with control cells (Figure 7E, F). These results suggest that, although Solo depletion increases radial keratin spoke density and partially reduces PG localization at cell-cell adhesion sites, it does not cause a detectable decrease in the mechanical strength of epithelial cell layers under the conditions examined.

**Figure 7.**
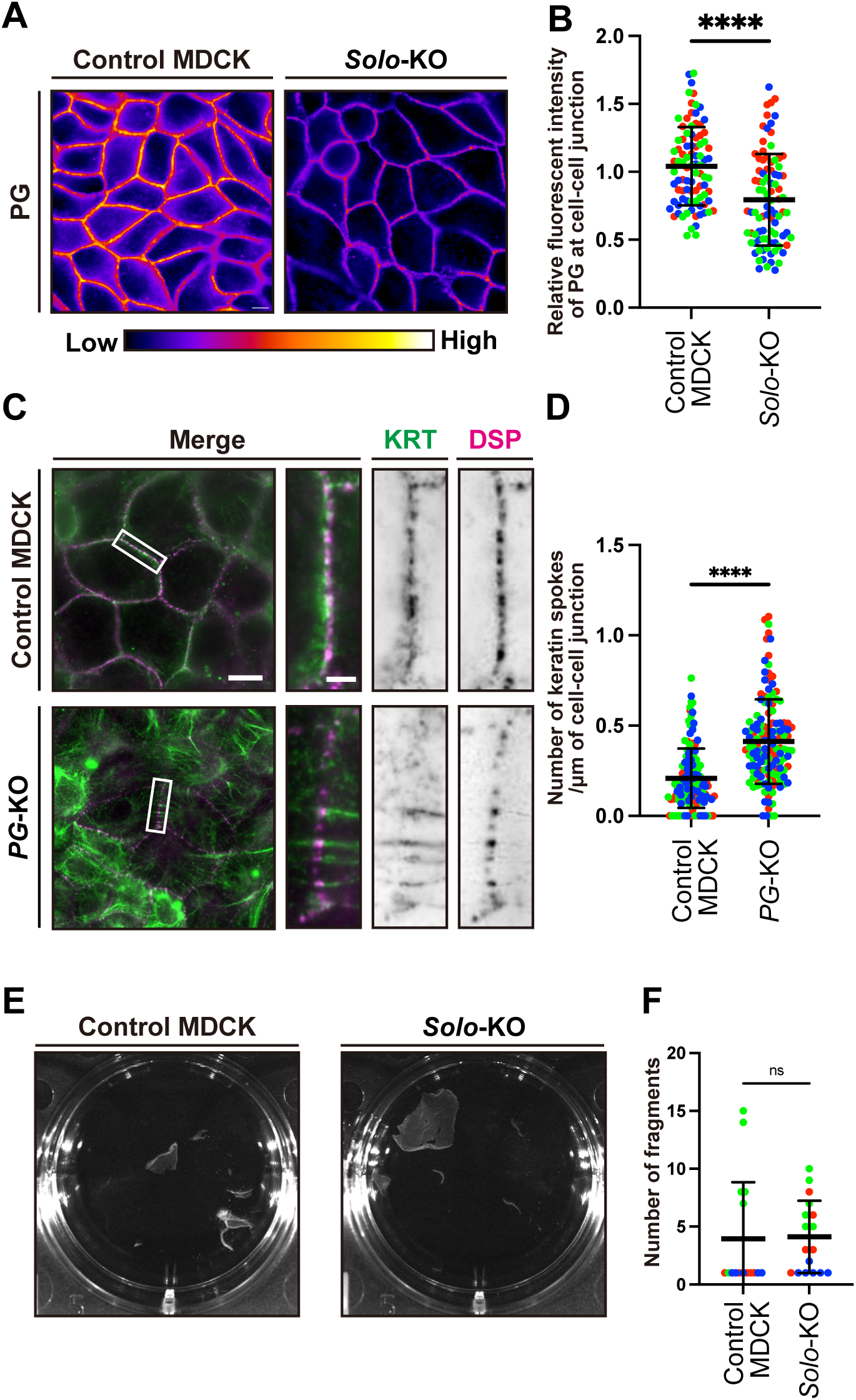
Solo facilitates PG localization to cell-cell adhesion sites and PG contributes to the reduction of radial keratin spoke density under confluent conditions. (A) Representative images showing PG localization in control and *Solo*-KO cells cultured under confluent conditions. Cells were fixed and stained with anti-PG antibodies. Scale bars, 10 µm. (B) Quantitative analysis of PG intensity at cell-cell adhesion sites shown in (A). Data are presented as the mean ± SD from three independent experiments, with 30 cell-cell junctions analyzed per experiment. ****, *p* < 0.0001 (Student’s t-test). (C) Representative images of keratin networks in control and *PG*-KO MDCK cells cultured under confluent conditions. Cells were fixed and stained with anti-pan-keratin (green) and anti-desmoplakin (DSP; magenta) antibodies. (D) Quantitative analysis of the density of spokes at cell-cell adhesion sites shown in (C). Data are presented as the mean ± SD from three independent experiments, with 42-55 cell-cell junctions analyzed per experiment. ****, *p* < 0.0001 (Student’s t-test) (E) Representative photographs of the results of the dispase assay. Monolayer cell sheets of control and *Solo*-KO cells were detached by dispase treatment and subjected to mechanical stress by vortexing for 30 s to assess cell sheet strength. (F) Quantitative analysis of the number of cell sheet fragments shown in (E). Data are presented as the mean ± SD from three independent experiments, with 5-6 cell sheets analyzed per experiment. ns, not significant (Student’s t-test)

## Discussion

In this study, we identified Solo as a regulator of cell density-dependent remodeling of keratin IFs at cell-cell adhesion sites. Increasing cell density was associated with a reduction in radial keratin spoke density and an increase in DSM length, consistent with DSM maturation. These changes were impaired in *Solo*-KO cells, which exhibited increased radial keratin spoke density and shorter DSMs under confluent conditions. The reduction in radial keratin spokes required the GEF activity of Solo and was sensitive to inhibition of actin polymerization and ROCK activity. In addition, loss of Solo partially reduced PG localization at cell-cell adhesion sites, whereas loss of PG increased radial spoke density. Together, these findings suggest that Solo contributes to cell density-dependent remodeling of DSM-associated keratin IFs and DSM maturation, potentially through Rho-ROCK-dependent actin cytoskeletal remodeling and the regulation of PG localization.

The reduction in radial keratin spokes appears to be associated with DSM maturation. During the early stages of cell-cell adhesion formation, DSMs are immature, and desmosomal cadherins initially form Ca²⁺-dependent interactions with cadherins on adjacent cells (Garrod and Chidgey, 2008; Watt *et al*., 1984). As cell density increases and cells remain in culture, these interactions become Ca²⁺-independent, resulting in the formation of mature DSMs that represent the steady-state form of epithelial cell-cell adhesion (Mattey and Garrod, 1986). In MDCK cells, conversion to the Ca²⁺-independent state requires high cell density and several days of culture (Mattey and Garrod, 1986). In our Ca^2+^-switch experiments, radial keratin spoke density initially increased after cell-cell contacts were re-established and subsequently decreased under confluent conditions. This temporal pattern suggests that spoke reduction may represent an early structural event associated with DSM maturation. Consistent with this interpretation, DSMs became longer along cell-cell junctions under confluent conditions, whereas this increase was impaired in *Solo*-KO cells. Thus, the transition from a radial spoke-rich organization toward a more rim-like keratin filament organization may accompany DSM maturation and enlargement. However, further studies are needed to determine whether these changes are mechanistically coupled or represent parallel aspects of junctional remodeling.

How Solo regulates this process remains unclear, but our findings implicate Solo-dependent actin cytoskeletal remodeling. Solo localization at cell-cell adhesion sites increased under confluent conditions, and its GEF activity was required for the reduction in radial keratin spoke density. Inhibition of actin polymerization or ROCK activity suppressed this reduction, suggesting that Rho-ROCK-dependent actin cytoskeletal remodeling contributes to the regulation of radial keratin spokes. In contrast, blebbistatin had no significant effect, suggesting that myosin II-dependent contractility may not be essential for this process. However, because blebbistatin was used at a concentration that preserved cell morphology, incomplete inhibition of myosin II activity cannot be excluded. Thus, actin cytoskeletal remodeling downstream of Rho-ROCK may be more important than myosin II-dependent contractility per se for the cell density-dependent remodeling of keratin–DSM structures. We previously demonstrated that Solo participates in cellular responses to tensile forces and that its interaction with keratin filaments is required for this response (Abiko et al., 2015; Fujiwara et al., 2016). These findings raise the possibility that changes in the mechanical environment associated with increasing cell density regulate Solo localization or activity at cell-cell adhesion sites. In this model, Solo-dependent RhoA signaling promotes actin remodeling, thereby contributing to DSM maturation and the associated reorganization of keratin IFs.

PG may be an important downstream component of this process. We previously showed that Solo knockdown reduces PG localization at cell-cell adhesion sites (Isozaki et al., 2020), and we observed a similar, albeit partial, reduction in PG localization in *Solo*-KO cells. Conversely, *PG*-KO cells exhibited a marked increase in radial keratin spoke density. Because Solo and PG do not exhibit identical localization patterns, PG recruitment to cell-cell adhesion sites is unlikely to be mediated solely by direct binding to Solo. Instead, Solo may regulate PG localization indirectly through actin cytoskeletal remodeling. The partial reduction in PG localization in *Solo*-KO cells further suggests that other factors contribute to PG localization or retention at these sites. Thus, Solo-dependent actin remodeling may facilitate PG accumulation at DSMs, which may, in turn, contribute to DSM maturation and the transition from radial keratin spokes toward rim-like keratin organization.

The relationship between keratin organization and epithelial mechanical strength appears to be complex. *Plectin*-KO cells exhibit increased radial keratin spoke density and reduced rim formation, accompanied by a modest reduction in cell-sheet strength (Prechova et al., 2022). In contrast, *Solo*-KO cells showed no significant reduction in cell layer strength despite pronounced defects in radial spoke remodeling and DSM maturation. Loss of PG, however, has been reported to substantially weaken epithelial cell adhesion (Yin et al., 2005). These observations suggest that changes in DSM-associated keratin IF organization are not necessarily accompanied by corresponding changes in the overall mechanical strength of the epithelial cell layer. Solo may primarily regulate the organization and maturation of DSM-associated structures, whereas other DSM components, including PG, may make distinct contributions to maintaining the mechanical integrity of cell-cell adhesions. Thus, cell density-dependent remodeling of radial spokes may provide a means of reorganizing keratin–DSM structures without necessarily causing a detectable change in the overall mechanical strength of the epithelial sheet.

Our findings also provide insight to the RhoGEF network that may act downstream of Solo. We previously demonstrated that Solo interacts with the RhoA-targeting GEFs LARG and PRG and that these interactions promote actin polymerization in response to substrate stiffness (Kunitomi et al., 2024; Kunitomi et al., 2026). In the present study, both *LARG*-KO and *PRG*-KO cells exhibited increased radial spoke density under confluent conditions, although the increases were less pronounced than that observed in *Solo*-KO cells. These findings suggest that LARG and PRG contribute to the regulation of radial keratin spoke density downstream of Solo and raise the possibility that they have partially redundant roles in RhoA signaling at cell-cell adhesion sites. However, functional redundancy has not been directly demonstrated, and whether Solo regulates the activity or localization of LARG and PRG at DSMs remains unknown. Further studies are needed to determine how cell density and mechanical cues regulate Solo at cell-cell adhesion sites and how the Solo-dependent RhoA–actin signaling is linked to PG localization and DSM maturation.

Taken together, our findings support a model in which Solo acts as a mechanochemical regulator linking changes in the cellular environment to the remodeling of keratin–DSM structures. Increasing cell density is associated with increased Solo localization at cell-cell adhesion sites, where Solo GEF activity and actin cytoskeletal remodeling contribute to the reduction of radial keratin spokes. Solo may also facilitate PG localization, thereby contributing to DSM maturation and enlargement. These changes are accompanied by a shift in keratin IF organization from radial keratin spokes toward rim-like structures connecting neighboring DSMs. This remodeling may enable epithelial cells to adapt the organization of cell-cell adhesion structures to changes in their mechanical environment without necessarily compromising overall tissue integrity. Further studies to define the molecular mechanisms regulating Solo localization and activity, and to determine how Solo-dependent signaling is coordinated with keratin and DSM remodeling, will be important for better understanding how epithelial tissues adapt their cytoskeletal architecture to changes in cell density and mechanical stress.

## Acknowledgments

We thank Kazuhisa Nakayama (Kumamoto University), Yohei Katoh (Hiroshima University) for technical advice.

## Author Declaration

### Data Availability Statement

Data that support the results reported here are available from the corresponding author, K.O. and S.C., upon reasonable request.

### Author Contribution Statement

Conceptualization: KM, KO, SC.

Funding acquisition: KO., SC.

Investigation: RM, UK.

Supervision: KO, SC.

Visualization: RM, UK.

Writing – original draft: KO, SC, UK.

Writing – review & editing: KO, SC, UK, KM.

### Ethics Approval and Consent to Participate

Not applicable.

### Patient Consent for Publication

Not applicable.

## ABBREVIATION

AJ: aadherens junction,
DSM: aDesmosome,
DSP: aDesmoplakin,
ExM: aexpansion microscopy,
IF: aintermediate filament,
KRT8: akeratin 8,
KRT18: akeratin 18,
KD: aknockdown,
KO: aknockout,
LINC: alinker of nucleoskeleton and cytoskeleton,
MDCK: aMadin-Darby canine kidney,
PG: aPlakoglobin,
PRG: aPDZ-RhoGEF,
RhoGEF: aRho-guanine nucleotide exchange factor,
ROCK: aRho-associated coiled-coil-containing protein kinase,
TJ: atight junction

