## Supplemental figures for "RhoGEF Solo regulates cell density-dependent keratin remodeling and desmosome maturation"

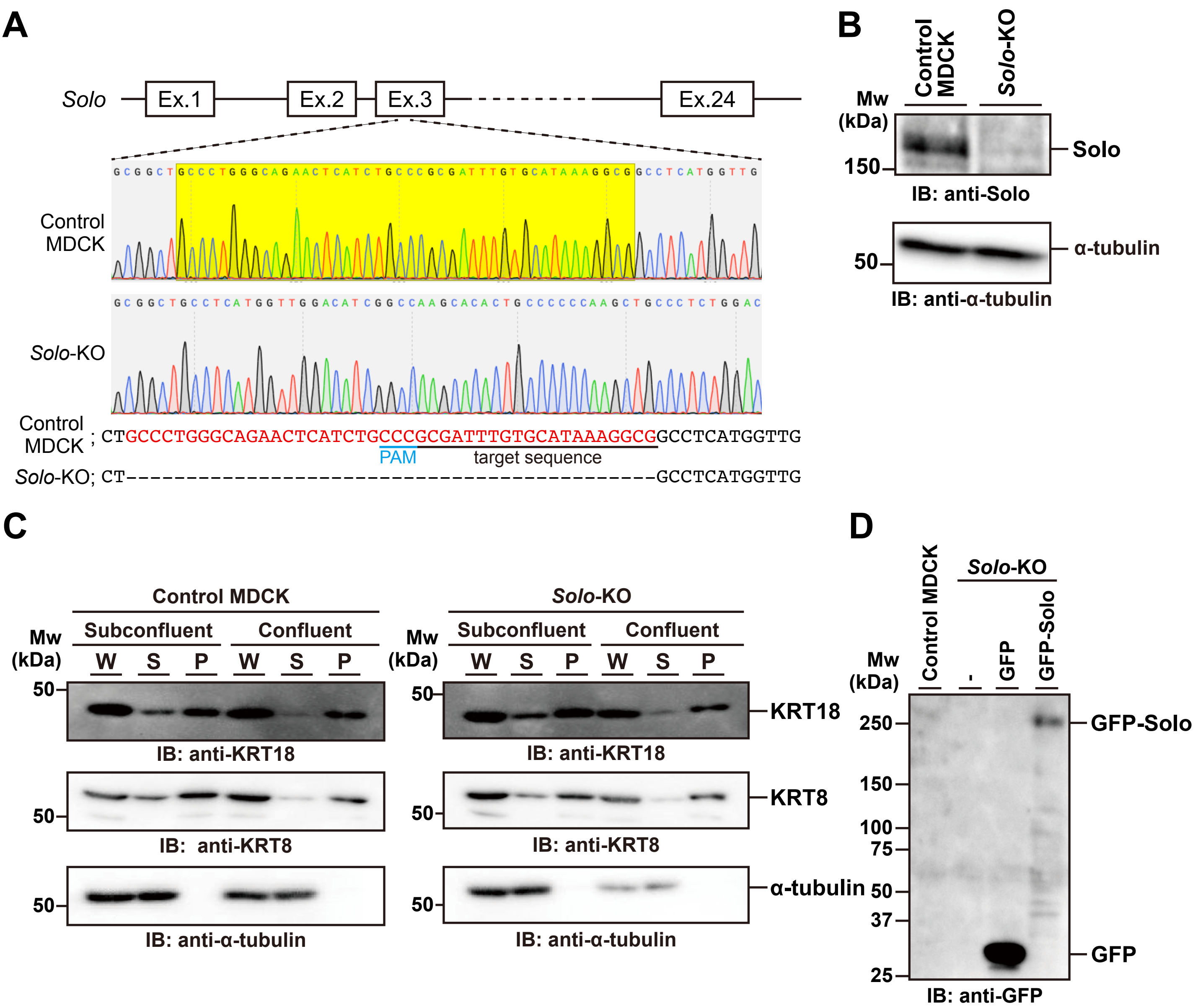


**Supplemental figure S1**

**Generation and characterization of a Solo-KO cell line from MDCK cells using CRISPR/Cas9 genome editing.**

(A) A schematic representation of the Solo gene locus and Sanger sequencing chromatograms of genomic DNA from control and Solo-KO cells. The target and PAM sequences are underlined. The deleted sequence in Solo-KO cells is highlighted in yellow and indicated in red characters. A 44-bp deletion was introduced into exon 3 of the Solo gene. (B) Immunoblots showing the loss of Solo protein expression in Solo-KO cells. (C) The soluble-to-insoluble ratios of KRT8 and KRT18 in control and Solo-KO cells. Whole cell lysates (W), Triton X-100-soluble (S), and insoluble (P) fractions were prepared from control and Solo-KO cells under subconfluent and confluent conditions. Immunoblot analysis was performed using antibodies against KRT8, KRT18, and α-tubulin. (D) Expression of EGFP and EGFP-Solo in Solo-KO cells was detected by western blotting using an anti-GFP antibody. In (B) and (C), α-tubulin was used as an internal loading control.

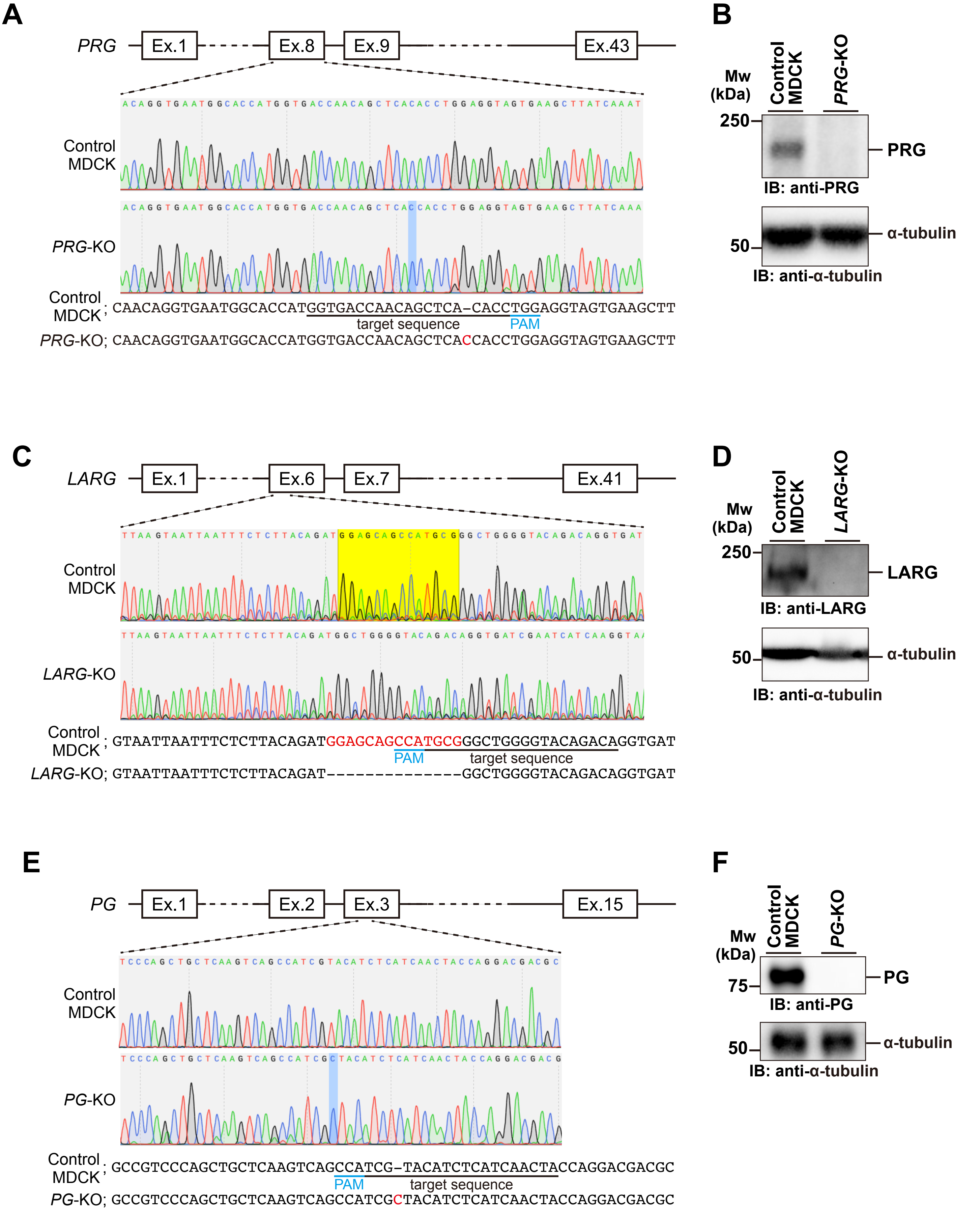


**Supplemental figure S2**

**Establishment of the PRG-KO, LARG-KO, and PG-KO cells from MDCK cells by CRISPR/Cas9 genome editing.**

(A) Schematic of the PRG gene locus and Sanger sequencing chromatograms of genomic DNA from control and PRG-KO cells. A 1-bp insertion was introduced into exon 8 of the PRG gene by a CRISPR/Cas9-mediated double-strand break. (B) Immunoblots showing the loss of PRG protein expression in PRG-KO cells. (C) Schematic of the LARG gene locus and Sanger sequencing chromatograms of genomic DNA from control and LARG-KO cells. A 14-bp deletion was introduced into exon 6 of the LARG gene. (D) Immunoblots showing the loss of LARG protein expression in LARG-KO cells. (E) Schematic of the PG gene locus and Sanger sequencing chromatograms of genomic DNA from control and PG-KO cells. A 1-bp insertion was introduced into exon 3 of the PG gene. (F) Immunoblots showing the loss of PG protein expression in PG-KO cells. (B, D, and F) α-tubulin was used as an internal loading control.
